# Computational Pan-Genomics: Status, Promises and Challenges

**DOI:** 10.1101/043430

**Authors:** The Computational Pan-Genomics Consortium, Tobias Marschall, Manja Marz, Thomas Abeel, Louis Dijkstra, Bas E. Dutilh, Ali Ghaffaari, Paul Kersey, Wigard P. Kloosterman, Veli Mäkinen, Adam M. Novak, Benedict Paten, David Porubsky, Eric Rivals, Can Alkan, Jasmijn Baaijens, Paul I. W. De Bakker, Valentina Boeva, Raoul J. P. Bonnal, Francesca Chiaromonte, Rayan Chikhi, Francesca D. Ciccarelli, Robin Cijvat, Erwin Datema, Cornelia M. Van Duijn, Evan E. Eichler, Corinna Ernst, Eleazar Eskin, Erik Garrison, Mohammed El-Kebir, Gunnar W. Klau, Jan O. Korbel, Eric-Wubbo Lameijer, Benjamin Langmead, Marcel Martin, Paul Medvedev, John C. Mu, Pieter Neerincx, Klaasjan Ouwens, Pierre Peterlongo, Nadia Pisanti, Sven Rahmann, Ben Raphael, Knut Reinert, Dick de Ridder, Jeroen de Ridder, Matthias Schlesner, Ole Schulz-Trieglaff, Ashley D. Sanders, Siavash Sheikhizadeh, Carl Shneider, Sandra Smit, Daniel Valenzuela, Jiayin Wang, Lodewyk Wessels, Ying Zhang, Victor Guryev, Fabio Vandin, Kai Ye, Alexander Schönhuth

**Affiliations:** Center for Bioinformatics, Saarland University, Saarbrücken, Germany; Max Planck Institute for Informatics, Saarbrücken, Germany; Bioinformatics and High Throughput Analysis, Faculty of Mathematics and Computer Science, Friedrich Schiller University Jena, Leutragraben 1, 07743 Jena, Germany; FLI Leibniz Institute for Age Research, Beutenbergstraße 11, 07745 Jena, Germany; Michael Stifel Center Jena, Ernst-Abbe-Platz 2, 07743 Jena, Germany; German Centre for Integrative Biodiversity Research (iDiv) Halle-Jena-Leipzig; The Delft Bioinformatics Lab, Department of Intelligent Systems, Delft University of Technology, Mekelweg 4, 2628 CD, Delft, The Netherlands; Istituto Nazionale Genetica Molecolare INGM, ‘Romeo ed Enrica Invernizzi’, Milan 20122, Italy.; Computational Science Lab, University of Amsterdam, Amsterdam, 1098XG, The Netherlands; Department of High Performance Computing, ITMO University, Saint Petersburg, 197101, Russia; Radboud Institute for Molecular Life Sciences, Center for Molecular and Biomolecular Informatics, Radboud University Medical Center, Nijmegen, Netherlands; Theoretical Biology and Bioinformatics, Utrecht University, Utrecht, Netherlands; Department of Marine Biology, Institute of Biology, Federal University of Rio de Janeiro, Rio de Janeiro, Brazil; EMBL-European Bioinformatics Institute, Wellcome Trust Genome Campus, Hinxton, CB10 1SD, UK; Department of Genetics, Center for Molecular Medicine, University Medical Center Utrecht, 3584 CG, Utrecht, The Netherlands; HIIT and Department of Computer Science, University of Helsinki, Finland; Genomics Institute, University of California Santa Cruz, Santa Cruz, CA 95064, USA; European Research Institute for the Biology of Ageing, University Medical Center Groningen, University of Groningen, Antonius Deusinglaan 1, AV Groningen 9713, The Netherlands; LIRMM, CNRS and Université de Montpellier, Montpellier, France; Institut de Biologie Computationnelle, CNRS and Université de Montpellier, Montpellier, France; Department of Computer Engineering, Bilkent University, Bilkent, Ankara, 06800, Turkey; Life Sciences Group, Centrum Wiskunde & Informatica, Amsterdam, 1098XG, The Netherlands; Institut Curie, Centre de Recherche, Inserm U900, F-75005 Paris, France; Mines ParisTech, F-77305 cedex Fontainebleau, France; PSL Research University, F-75005 Paris, France; Institut Cochin, Inserm U1016, CNRS UMR 8104, Université Paris Descartes UMR-S1016, F-75014 Paris, France; Departments of Statistics, The Pennsylvania State University, University Park, PA, 16802; CNRS, Univ. Lille, UMR 9189 CRIStAL, F-59000 Lille, France; Division of Cancer Studies, King’s College London, London SE11UL, UK; MonetDB Solutions, Amsterdam, The Netherlands; KeyGene N.V, Agro Business Park 90, 6708 PW Wageningen, The Netherlands; Department of Epidemiology, Erasmus Medical Center, Rotterdam, The Netherlands; Department of Genome Sciences, University of Washington, Seattle, WA, 98195, USA; Howard Hughes Medical Institute, University of Washington, Seattle, WA, 98195, USA; Genome Informatics, Institute of Human Genetics, University Hospital Essen, University of Duisburg-Essen, Essen, Germany; Department of Computer Science, University of California, Los Angeles, USA; Department of Human Genetics, University of California, Los Angeles, USA; Wellcome Trust Sanger Institute, Cambridge, UK; Centre for Integrative Bioinformatics VU (IBIVU), VU University Amsterdam, De Boelelaan 1081A, 1081 HV Amsterdam, The Netherlands; Center for Computational Molecular Biology and Department of Computer Science, Brown University, Providence, RI 02912, USA; European Molecular Biology Laboratory (EMBL), Genome Biology Unit, Meyerhofstrasse 1, 69117 Heidelberg, Germany; Genomics Coordination Center, University of Groningen, University Medical Center Groningen, Groningen, 9700RB, The Netherlands; Department of Computer Science and Center for Computational Biology, Johns Hopkins University, Baltimore, Maryland; Science for Life Laboratory, Dept. of Biochemistry and Biophysics, Stockholm University, Box 1031, SE-17121 Solna, Sweden; Department of Computer Science and Engineering, The Pennsylvania State University, USA; Department of Biochemistry and Molecular Biology, The Pennsylvania State University, USA; Genomic Sciences Institute of the Huck, The Pennsylvania State University, USA; Bina Technologies, Roche Sequencing, Redwood City, CA 94065, USA; Biological Psychology, Vrije Universiteit Amsterdam, The Netherlands; Genalice BV, Harderwijk, The Netherlands; INRIA Campus de Beaulieu-Rennes, Rennes Cedex 35042, France; Dipartimento di Informatica, University degli Studi di Pisa, Pisa, Italy; Erable Team, INRIA; Department of Mathematics and Computer Science, Freie Universität Berlin, Berlin, Germany; Bioinformatics Group, Wageningen University, Droevendaalsesteeg 1, 6708 PB, Wageningen, The Netherlands; Division of Theoretical Bioinformatics, German Cancer Research Center (DKFZ), Im Neuenheimer Feld 280, 69120 Heidelberg, Germany; Illumina Cambridge Ltd, Chesterford Research Park, Little Chesterford, Essex CB10 1XL, UK; Terry Fox Laboratory, BC Cancer Agency, Vancouver, British Columbia, Canada; Leiden Observatory, Leiden University, P.O. Box 9513, 2300 RA Leiden, The Netherlands; School of Management, Xi’an Jiaotong University, Xi’an, Shaanxi 710049, China; Institute of Data Science and Information Quality, Xi’an Jiaotong University; Shaanxi Engineering Research Center of Medical and Health Big Data, Xi’an Jiaotong University; Netherlands Cancer Institute (NKI), Amsterdam, The Netherlands; Department of Mathematics and Computer Science, University of Southern Denmark, Campusvej 55, DK-5230, Odense M, Denmark; School of Electronic and Information Engineering, Xi’an Jiaotong University, Xi’an, Shaanxi 710049, China; Kai Ye Young Scientist Studio, Xi’an Jiaotong University

## Abstract

Many disciplines, from human genetics and oncology to plant breeding, microbiology and virology, commonly face the challenge of analyzing rapidly increasing numbers of genomes. In case of *Homo sapiens*, the number of sequenced genomes will approach hundreds of thousands in the next few years. Simply scaling up established bioinformatics pipelines will not be sufficient for leveraging the full potential of such rich genomic datasets. Instead, novel, qualitatively different computational methods and paradigms are needed. We will witness the rapid extension of *computational pan-genomics*, a new sub-area of research in computational biology. In this paper, we generalize existing definitions and understand a *pan-genome* as any collection of genomic sequences to be analyzed jointly or to be used as a reference. We examine already available approaches to construct and use pan-genomes, discuss the potential benefits of future technologies and methodologies, and review open challenges from the vantage point of the above-mentioned biological disciplines. As a prominent example for a computational paradigm shift, we particularly highlight the transition from the representation of reference genomes as strings to representations as graphs. We outline how this and other challenges from different application domains translate into common computational problems, point out relevant bioinformatics techniques and identify open problems in computer science. With this review, we aim to increase awareness that a joint approach to computational pan-genomics can help address many of the problems currently faced in various domains.

## 1 Introduction

In 1995, the complete genome sequence for the bacterium *Haemophilus influenzae* was published [1], followed by the sequence for the eukaryote *Saccharomyces cerevisiae* in 1996 [2] and the landmark publication of the human genome in 2001 [3, 4]. These sequences, and many more that followed, have served as *reference genomes*, which formed the basis for both major advances in functional genomics and for studying genetic variation by re-sequencing other individuals from the same species [5, 6, 7, 8]. The advent of rapid and cheap “next-generation” sequencing technologies since 2006 has turned re-sequencing into one of the most popular modern genome analysis workflows. As of today, an incredible wealth of genomic variation within populations has already been detected, permitting functional annotation of many such variants, and it is reasonable to expect that this is only the beginning.

With the number of sequenced genomes steadily increasing, it makes sense to re-think the idea of a *reference* genome [9, 10]. Such a reference sequence can take a number of forms, including:

- the genome of a single selected individual,
- a consensus drawn from an entire population,
- a “functional” genome (without disabling mutations in any genes), or
- a maximal genome that captures all sequence ever detected.

Depending on the context, each of these alternatives may make sense. However, many early reference sequences did not represent any of the above. Instead, they consisted of collections of sequence patches, assayed from whatever experimental material had been available, often from a relatively unstructured mix of individual biological sources. Only lately has the rapid spread of advanced sequencing technologies allowed the reasonably complete determination of many individual genome sequences from particular populations [7], taxonomic units [11], or environments [12]. To take full advantage of these data, a good “reference genome” should have capabilities beyond the alternatives listed above. This entails a paradigm shift, from focusing on a single reference genome to using a *pan-genome*, that is, a representation of all genomic content in a certain species or phylogenetic clade.

### 1.1 Definition of Computational Pan-Genomics

The term *pan-genome* was first used by Sigaux [13] to describe a public database containing an assessment of genome and transcriptome alterations in major types of tumors, tissues, and experimental models. Later, Tettelin et al. [9] defined a microbial pan-genome as the combination of a *core* genome, containing genes present in all strains, and a *dispensable* genome (also known as flexible or accessory genome) composed of genes absent from one or more of the strains. A generalization of such a representation could contain not only the genes, but also other variations present in the collection of genomes. The idea of transitioning to a human pangenome is also gaining attention^1^. While the pangenome is thus an established concept, its (computational) analysis is frequently still done in an ad-hoc manner.

Here, we generalize the above definitions and use the term *pan-genome* to refer to any collection of genomic sequences to be analyzed jointly or to be used as a reference. These sequences can be linked in a graph-like structure, or simply constitute sets of (aligned or unaligned) sequences. Questions about efficient data structures, algorithms and statistical methods to perform bioinformatic analyses of pan-genomes give rise to the discipline of *computational pan-genomics*.

Our definition of a pan-genome allows to capture a diverse set of applications across different disciplines. Examples include “classical” pan-genomes consisting of sets of genes present in a species [14], graph-based data structures used as references to enhance the analysis of difficult genomic regions such as the MHC [15], the compact representation of a transcriptome [16], and the collection of virus haplotypes found in a single patient [17].

While being aware that the above definition of a pan-genome is general, we argue that it is instrumental for identifying common computational problems that occur in different disciplines. Our notion of computational pan-genomics therefore intentionally intersects with many other bioinformatics disciplines. In particular it is related to *metagenomics*, which studies the entirety of genetic material sampled from an environment; to *comparative genomics*, which is concerned with retracing evolution by analyzing genome sequences; and to *population genetics*, whose main subject is the change of a population’s genetic composition in response to various evolutionary forces and migration. While none of these fields captures all aspects of pangenomics, they all have developed their own algorithms and data structures to represent sets of genomes and can therefore contribute to the pangenomics toolbox. By advocating *computational pan-genomics*, we hope to increase awareness of common challenges and to generate synergy among the involved fields.

At the core of pan-genomics is the idea of replacing traditional, linear reference genomes by richer data structures. The paradigm of a single reference genome has endured in part because of its simplicity. It has provided an easy framework within which to organize and think about genomic data; for example, it can be visualized as nothing more than linear text, which has allowed the development of rich two-dimensional genome browsers [18, 19]. With the currently rapidly growing number of sequences we have at our disposal, this approach increasingly fails to fully capture the information on variation, similarity, frequency, and functional content implicit in the data. Although pan-genomes promise to be able to represent this information, there is not yet a conceptual framework or a toolset for working with pangenomes that has achieved widespread acceptance. For many biological questions, it is not yet established how to best extract the relevant information from any particular pan-genome representation, and even when the right approach can be identified, novel bioinformatics tools often need to be developed in order to apply it.

In this paper, we explore the challenges of working with pan-genomes, and identify conceptual and technical approaches that may allow us to organize such data to facilitate its application in (green, blue, red, and white [20]) biotechnology and fundamental research.

### 1.2 Goals of Computational PanGenomics

On a high level, desirable features of a pan-genome include *completeness*, or containing all functional elements and enough of the sequence space to serve as a reference for the analysis of additional individuals;*stability*, or having uniquely identifiable features that can be studied by different researchers and at different points in time;*comprehensibility*, or facilitating understanding of the complexities of genome structures across many individuals or species; and *efficiency*, or organizing data in such a way as to accelerate downstream analysis.

These desiderata highlight the breadth of challenges facing pan-genomics as a field, some of which go beyond scientific questions. Reaching *completeness*, for instance, requires the necessary (financial and technical) resources to collect and sequence a sufficient number of genomes for a particular tissue, organism, species, other taxonomic unit, ecological community, or geospatial niche of interest to be accurately represented. The availability of data sharing mechanisms will greatly influence how quickly *completeness* can be achieved. Issues of data sharing include technical ones (mostly due to the data being big), political ones, and ethical/privacy concerns [21], as well as issues related to the interplay of these three areas. Achieving *stability* requires a central, recognized authority equipped with the long-term resources for curating reference pan-genomes. Besides this organizational component, achieving stability also requires reaching consensus about ways to define coordinate systems on pan-genomes. The goal of *comprehensibility* is mostly a biological problem. What it means exactly can differ substantially between application domains. The goal of *efficiency*, on the other hand, is in the domain of computer science. Aligning the needs of researchers in the application domains with efforts to develop algorithms and statistical methods is key to designing efficient solutions. With this paper, we hope to contribute significantly to this communication process.

## 2 Applications

Pan-genomes arise in many different application domains. In this section, we discuss seven different fields: Microbes, Metagenomics, Viruses, Plants, Human Genetic Diseases, Cancer, and Phylogenomics. While this list of areas related to pangenomics is not exhaustive, we aimed to select those that we believe are most strongly impacted by novel pan-genomics methods, and hence are most relevant to our goal of identifying common computational challenges.

### 2.1 Microbes

Bacteria and fungi are widely studied—and applied—in fields including biology, medicine, and biotechnology. A full understanding of the functional and evolutionary repertoire of microbial genomes thus not only is interesting from a scientific point of view, but also opens up possibilities for developing therapies and engineering applications.

For a number of microorganisms, pan-genome sequence data is already available; refer to [22, 23] for examples. Microbes provide a unique opportunity for pan-genome construction: the size of their genomes is relatively small, and for many species there are multiple fully closed genome sequences available. Furthermore, for some clinically interesting bacterial species, up to thousands of sequenced strains are available at sufficient depth to create draft genome assemblies. This has enabled pangenome studies at the gene level [14], for which established workflows and mature software are available, as reviewed in [24]. With the current data, however, we are in a position to create a pangenome at the sequence level, as in e.g. [25]. In this context, a pan-genome is a representation that encodes the complete sequence information of many individual strains.

From an evolutionary point of view, microbial pan-genomes support comparative genomics studies. These are particularly interesting due to most microorganisms’ potential for horizontal gene exchange. This means that not all genes in a genome adhere to the same phylogenetic sub-tree [26]. Thus, the evolution of microorganisms including bacteria, but also higher organisms [27], is more naturally represented as a phylogenetic network, rather than a phylogenetic tree [28]. We envision that these phylogenetic networks can be encoded in the structure of the pan-genome.

Applying genome-wide association studies (GWAS) to microbes is an emerging field [29, 30], promising to pinpoint genetic variables that correlate with relevant traits such as drug resistance or secondary metabolism. Such studies can operate at the level of individual variants—such as single-nucleotide polymorphisms (SNPs), insertion/deletion variants (indels), and structural variants (SVs)—or at the level of absence or presence of whole genes, annotated functions, or mobile genetic elements such as integrons or prophages. Computational pan-genomic approaches could be applied at each of these levels. Important challenges amenable to a pan-genomic approach include establishing reliable data processing pipelines to deliver variant calls, extracting gene absence/presence signals from NGS data, annotation for hypothetical genes and proteins, and specifically computational challenges such as the definition of a coordinate system to identify sequence loci on pan-genomes or the handling nested variation, such as SNP positions in large insertions. By addressing these challenges, computational pan-genomics has the potential to substantially contribute to the success of microbial GWAS.

### 2.2 Metagenomics

Metagenomics studies the genomic composition of microorganisms sampled from an environment. Abundant metagenomic data is currently being generated from various environments such as human hosts [12, 31], the world’s oceans [32, 33], and soil [34]. One main advantage of this approach lies in allowing the sampling of *all* microorganisms in an environment, not only those that can be cultured. This however comes at the cost of having to untangle the sequencing data generated from such a mixture computationally. A first question often asked is about the taxonomic composition of the sample. Other relevant questions that can be approached with metagenomic data include ascertaining the presence of certain gene products or whole pathways, and determining which genomes these functional genes are associated with.

Metagenomics can be applied to gain insights on human health and disease. Metagenome-wide association studies that aim to associate the microbial composition in the human gut with diseases such as type 2 diabetes are an example [35]. Metagenomics has also been shown to be capable of revealing the genomes of entire species, and tracing them through environments, as in the example of the shiga-toxigenic Escherichia coli being responsible for a recent major outbreak in Germany [36].

In the metagenomic setting, the set of genomic sequences underlying a pan-genome is not defined by ancestral relationships, but by co-occurrence in an environment. This presents both a challenge and an opportunity. On the one hand, constructing such a pan-genome and drawing robust conclusions from it is difficult, especially when sequencing reads are short. On the other hand, it presents the chance to reveal common adaptations to the environment as well as co-evolution of interactions.

### 2.3 Viruses

Viruses are notorious mutation machines. A viral quasi-species is a cloud of viral haplotypes that surround a given master virus [17]. Although viral genomes are comparatively short (RNA viruses range from 3–30kb, DNA viruses are usually not larger than 3Mb), their high sequence variability makes it challenging to assemble full viral genomes *de novo*. There are two major sequencing approaches for viruses: sequencing isolated viral clones, and metagenomic sequencing. The latter usually identifies a metapopulation consensus genome sequence rather than a single haplotype [37], and includes confounding genetic sequences such as the genome of other community members and of the cellular virus host. Thus far, the obvious approach of viral particle sorting by Fluorescence-Activated Cell Sorting (FACS), followed by single virus sequencing, has remained elusive due to their small genome size [38, 39]. New long-read technologies (e.g. PacBio, Oxford Nanopore) are now providing the first promising results in the sequencing of complete viral genomes [40, 41]. Currently, error rates in these third-generation long read sequencing technologies still far exceed the frequencies of rare strains or haplotypes. However, as sequencing chemistry and technologies progress, such techniques are likely to become key tools for the construction of viral pangenomes.

Low frequency strains are hardly detectable, especially for fast evolving RNA viruses with a replication mutation rate similar to sequencing error rates. Reliable viral haplotype reconstruction is not fully solved, although to date many promising approaches have been presented [42]. Haplotype resolution techniques such as Strand-Seq [43] are not applicable for small virus particles.

One of the goals of pan-genomics, both in virology and in medical microbiology, will be to fight infectious disease. We expect that computational pan-genomics will assist GWAS approaches, which may allow the prediction of crucial parameters such as the exact diagnosis, staging, and suitable therapy selection from a given patient’s viral pan-genome. For example, several studies have shown relationships between genetic diversity and disease progression, pathogenesis, immune escape, effective vaccine design, and drug resistance in HIV [44, 45, 46]. Thus, computational pangenomics promises to be useful when studying the response of the quasi-species to the host immune system, in the context of personalized medicine.

The molecular interactions between pathogens and their hosts lead to a genetic arms race that allows virus-host interactions to be predicted [47]. In this context, metagenomics techniques can also be applied [48, 49]. The large metagenomic datasets mentioned in Section 2.2 can serve as input for such studies. We expect that computational pangenomics will allow increased power and accuracy, for example by allowing the pan-genome structure of a viral population to be directly compared with that of a susceptible host population.

### 2.4 Plants

Genomic hybridization of accessions of crops or flowers has been exploited for over a century to create offspring with desirable traits. Genes found in wild varieties that improve important properties of crops, such as appearance, nutrient content, resistance to certain pests or diseases, or tolerance for stresses such as drought or heat, are now routinely bred into commercial crop varieties.

Large-scale genomics projects to characterize the genetic diversity in plants are already ongoing, not only for the model plant *Arabidopsis thaliana* [5], but also for crops [50]. Examples include the resequencing of hundreds to thousands of varieties of rice [51], maize [52], sorghum [53], and tomato [6]. Future projects aim to sequence many more varieties, e.g. 100,000 varieties of rice^2^. Mining and leveraging the sequence data in such large-scale projects requires a pan-genomic approach. Particularly challenging is the fact that many plant genomes are large, complex (containing many repeats) and often polyploid.

A pan-genome structure has multiple advantages over a single, linear reference genome sequence in plant breeding applications. Having a pan-genome available for a given crop that includes its wild relatives provides a single coordinate system to anchor all known variation and phenotype information, and will allow for identification of novel genes from the available germplasm that are not present in the reference genome(s). Moreover, the pangenome will reveal chromosomal rearrangements between genotypes which can hinder the introgression of desired genes. It also provides a compact representation of polyploid genomes and, in case of autopolyploids, allows for the quantitation of allele dosage between individuals.

### 2.5 Human Genetic Diseases

A key objective in human genetics is to develop a full understanding of how genotype influences phenotype. To date, numerous genes have been successfully mapped for rare monogenic diseases, where a rare mutation results, in most cases, in disease. For mutations with (almost) fully penetrant effects, linkage analysis in affected pedigrees has demonstrated to be a highly effective approach for localizing causal genes, with a few thousand of such disease genes now annotated in the Online Mendelian Inheritance in Man^3^. More recently, whole-exome sequencing in families offered a complimentary and perhaps more efficient approach, pinpointing culprit mutations in novel disease genes and in specific cases revealing a role for *de novo* mutations in affected offspring (without familial segregation). By comparing observed variants in affected family members to those present in the population at large (e.g., the Exome Variant Server (EVS)^4^ and the ExAC Browser^5^ [54]), it is possible to implicate such variants as being pathogenic on the basis of their frequency in the population. Yet caution is warranted, because any genome sequence may contain many potentially functional, rare variants that could result in false-positive claims about disease-causing variants [55].

Historically, common diseases have been more challenging to study, because they arise from the interplay of many genetic and non-genetic risk factors, each of which only contributing modestly to the overall risk. Thanks to systematic approaches to interrogate human genomic variation in large numbers, genome-wide association studies (GWAS) have identified thousands of robust genotype-phenotype associations for a wide range of human traits and diseases. Critical to this success have been catalogs of human genome variation, their linkage disequilibrium (LD) properties as well as the commercial development of cheap microarrays. One of the primary applications of the resources provided by the HapMap [56] and 1000 Genomes [8] (1KG) projects was LD-based imputation [57], allowing GWAS to test many more variants than typically present on a single microarray chip.

Although the 1KG database is predicted to include more than 99% of all variants with a frequency of at least 1% [8], many other independent sequencing efforts revealed that many variants below 1% remain to be discovered as we continue to sequence more and more samples. This is probably best illustrated by the Exome Aggregation Consortium database [54], which provides detailed information on 7.4 million high-quality variants in the exomes of 60,706 unrelated individuals. Indeed, 54% of the observed variants are singletons and an impressive 72% are not even observed in 1KG. This highlights that there will be enormous value for medical genetics in creating pan-genomic resources to aggregate common and rare variants for imputation and association testing of commonly segregating variants on the one hand, and the (functional) interpretation of rare variants in personal genome sequences on the other.

To the extent that common variants play an important role for certain traits, they are likely to be present in 1KG and can therefore in principle be imputed well and thus tested for association. This is, however, not the case for rare variants, which are poorly represented by 1KG, and challenging to impute from a microarray chip. We therefore see potential value in sequencing larger samples for diseases where common variants appear to contribute only modestly, as illustrated by the recent GWAS for amyotrophic lateral sclerosis (ALS) [58]. Ultimately, the key tradeoff in the “optimal” study design is the likelihood of finding novel discoveries (power) and the associated cost (efficiency). With sequencing still being a significant cost driver, there is no simple recommendation for diverse diseases.

In addition to the discovery of novel associations, we should also expect an increasing focus on the fine-mapping of the initial GWAS hits, that is, the localization of the causal variants driving the association signals. Identifying these variants (and their functional impact) is a key step in elucidating the biological mechanisms involved. Combining genome-wide variants with comprehensive functional annotations—for example, from epigenomic or gene expression data sets—should be considered a priority. We expect pan-genome data structures that are capable of handling such annotations to significantly contribute to this endeavor.

In parallel, we should expect substantial improvements in the characterization of parts of the genome that are currently not easily accessible with current sequencing technologies (including repetitive regions, or those of low complexity), and the detection of complex structural variants, as these are still more challenging to call from raw sequence data. Despite efforts to capture structural variation based on discordant mapping of short reads, a major fraction remains undetected in large part because of their complexity and due to the incompleteness of the current reference genome [59, 60]. Incorporating fully resolved high-quality structural variation data into a pan-genomic reference, preferably from long-read sequencing data, would greatly improve the genotyping of known structural variants and limit false-positives among novel variants. This will be highly relevant in the clinical setting, where genome sequencing is expected to replace array-based copy number variation profiling within a few years.

### 2.6 Cancer

Cancer is caused mostly by somatic DNA alterations that accumulate during an individual’s lifetime [61]. Somatic mutations in different individuals arise independently, and recent large cancer studies have uncovered extensive *inter-patient heterogeneity* among somatic mutations, with any two tumors presenting a different complement of hundreds to tens of thousands of somatic mutations [62, 63]. Heterogeneity also manifests *intra-patient*, with different populations of cells presenting different complements of mutations in the same tumor [64, 65].

Inter-patient and intra-patient heterogeneity pose several challenges to the detection and the interpretation of somatic mutations in cancer. The availability of a pan-genome reference would greatly improve the detection of somatic mutations in general, through improved quality of read mapping to polymorphic regions, and in particular in cases when matched normal tissue is not available or when only a reduced sequence coverage can be obtained.

In addition to a pan-genome reference, a somatic cancer pan-genome, representing the variability in the observed as well as inferred background alteration rate across the genome and for different cohorts of cancer patients, would enhance the identification of genomic alterations related to the disease (*driver events*) based on their recurrence across individuals. Even more important would be the availability of a somatic pan-genome describing the general somatic variability in the human population, which would provide an accurate baseline for assessing the impact of somatic alterations.

For the medium and long term future, we envision a comprehensive cancer pan-genome to be built for each tumor patient, comprising single-cell data, haplotype information as well as sequencing data from circulating tumor cells and DNA. Such a pan-genome will most likely constitute a much better basis for therapy decisions compared to current cancer genomes which mainly represent the most abundant cell type.

### 2.7 Phylogenomics

Phylogenomics reconstructs the evolutionary history of a group of species by using their complete genome sequences, and can exploit various signals such as sequence or gene content [66, 67]. Computational pan-genomics will allow genomic features with an evolutionary signal to be rapidly extracted, such as gene content tables, sequence alignments of shared marker genes, genome-wide SNPs, or internal transcribed spacer (ITS) sequences, depending on the level of relatedness of the included organisms. This will facilitate evolutionary analyses ranging from the reconstruction of species phylogenies, where heterogeneity between genomes is high [68], to tracing epidemic outbreaks and cancer lineages within a patient, where heterogeneity between genomes is low. For example, the yeast dataset described in [69] allowed a phylogenetic classification based on the presence and location of mobile elements in several strains of *S. cerevisiae*. Computational pan-genomics would also enhance such analyses when the pan-genome is built from a set of distinct strains of the same species.

Unambiguous phylogenomic trees of organismal or cellular lineages form invaluable input data for applications in various biomedical fields, for example to map the evolutionary dynamics of mutation patterns in genomes [70] or to understand the transfer of antibiotic resistance plasmids [71]. At the same time, the size of the pan-genome often hampers the inference of such a “tree of life” computationally as well as conceptually. One clear bonus offered by the pan-genome, is that for traditional phylogenomics only the best aligned, and most well behaved residues of a multiple sequence alignment can be retained. In contrast, the pangenomic representation of multiple genomes allows for a clear encoding of the various genomic mutations in a model of the evolutionary events. This leads to the possibility for radical new evolutionary discoveries in fields including the origin of complex life [72], the origin of animals [73] and plants [74], or the spread of pathogens [75, 76], but also inferring the relationships between cancer lineages within a single patient [77, 78].

## 3 Impact of Sequencing Technology on Pan-Genomics

Next-generation short-read sequencing has contributed tremendously to the increase in the known number of genetic variations in genomes of many species. The inherent limitations of commonly used short-read sequencing are three-fold. First, the short read lengths prohibit the interrogation of genomic regions consisting of repetitive stretches, the direct phasing of genetic variants [79], and the detection of large structural variations [60]. Second, non-random errors hamper the detection of genetic variations [80]. Third, there is a non-uniform distribution of sequencing coverage [81] due to various factors including biases in PCR amplification, polymerase processivity, and bridge amplification.

Establishing pan-genome sequences ideally requires a complete set of *phased*—that is, haplotype resolved—genetic variations. Experimental techniques to capture such linkage information have witnessed significant progress recently, as reviewed by [82]. Ultimately, specialized protocols for haplotype-resolved sequencing will be rendered obsolete once sufficiently long sequencing reads are routinely available.

The most promising developments in sequencing technology involve single-molecule real-time sequencing of native DNA strands. Currently, SMRT sequencing (Pacific Biosciences) is widely used for variation discovery and genome assembly [60]. The MinION device (Oxford Nanopore Technologies) [83] provides even longer reads of single DNA molecules, but has been reported to exhibit GC biases [84]. Data generated on the MinION platform has been successfully used for assembly of small genomes and for unraveling the structure of complex genomic regions [85, 86].

Despite this progress, sequencing reads are not yet sufficiently long to traverse and assemble all repeat structures and other complementary technologies are necessary to investigate large, more complex variation. Presently, array comparative genomic hybridization (arrayCGH), synthetic long reads (Moleculo [87], 10X Genomics [88]), chromatin interaction measurements [89] and high-throughput optical mapping [90, 91, 92] all aid the detection of structural variation.

Beyond interrogating genomes, sequencing technologies also serve to measure various other signals that can be seen as additional layers of information to be stored and analyzed in a pan-genome framework. Most notably, specialized protocols exist to measure transcriptomes, DNA-protein interaction, 3D genome structure, epigenetic information, or translation activity. In all these cases a current challenge consists in transitioning from bulk to single-cell sequencing.

We expect that novel technologies will continue to greatly improve all mentioned applications in genomics and beyond. Nonetheless, further decreasing costs and conducting appropriate benchmark studies that illustrate specificity and sensitivity are problems yet to be tackled.

## 4 Data Structures

Besides novel experimental protocols and sequencing technologies, advances in *data structures* play a key role for making pan-genome analyses a reality. In this section, we identify important design goals for pan-genome data structures and survey existing approaches.

First, we ground our discussion with a practical example of a pan-genome data structure. Figure 1 presents a splicing graph [16] for a single human gene. This compact representation of a collection of transcripts of a given gene has seen application in resequencing-based analyses, where it is used to support the alignment of RNA sequencing reads to the entire transcriptome [94]. It comprises genomic sequences, observed linkages between them, and the original transcripts and reference genome used to build the graph. On the one hand, this example illustrates that *computational pan-genomics* applies to collections of genetic sequences that are not necessarily whole genomes: in order to best support the intended application of analyzing RNA sequencing data, we here restrict our pan-genome data structure to transcribed sequences. On the other hand, this example highlights the importance of *graphs* for pan-genomic data structures. Here, the graph consists of sequences (nodes), of adjacencies between them (edges), and of the sequences that gave rise to it (paths). A wide array of pangenomic operations relies on the interplay between these basic elements.

### 4.1 Design Goals

Different applications give rise to different requirements for data structures that represent pangenomes. Figure 2 presents a schematic overview. Depending on the specific application, a pangenome data structure may need to offer any of the following capabilities:

#### Construction and Maintenance

Pangenomes should be constructable from different independent sources, such as (1) existing linear reference genomes and their variants, (2) haplotype reference panels, and (3) raw reads, either from bulk sequencing of complex mixtures or from multiple samples sequenced separately (see Figure 2, *Construct* operation). The data structure should allow dynamic updates of stored information without rebuilding the entire data structure, including local modifications such as adding a new genetic variant, insertions of new genomes and deletion of contained genomes (see Figure 2, box *Update* operation).

#### Coordinate System

A pan-genome defines the space in which (pan-)genomic analyses take place. It should provide a “coordinate system” to unambiguously identify genetic loci and (potentially nested) genetic variants. Desirable properties of such a “coordinate system” include that nearby positions should have similar coordinates, paths representing genomes should correspond to monotonic sequences of coordinates where possible, and coordinates should be concise and interpretable.

#### Biological Features and Computational Layers

Annotation of biological features should be coherently provided across all individual genomes (see Figure 2, *Annotate* operation). Computationally these features represent additional layers on top of pan-genomes. This includes information about (1) genes, introns, transcription factor binding sites; (2) epigenetic properties; (3) linkages, including haplotypes; (4) gene regulation; (5) transcriptional units; (6) genomic 3D structure and (7) taxonomy among individuals.

#### Data Retrieval

A pan-genome data structure should provide positional access to individual genome sequences, access to all variants and to the corresponding allele frequencies (see Figure 2, *Retrieve* operation). Haplotypes should be reconstructable including information about all maximal blocks and linkage disequilibrium between two variants.

#### Searching within Pan-Genomes

Comparisons of short and long sequences (e.g. reads) with the pan-genome ideally results in the corresponding location and the best matching individual genome(s) (see Figure 2, *Map* operation). This scenario may occur for transcriptomic data as well as for DNA re-sequencing data, facilitating the identification of known variants in new samples (see Figure 2, *Variant calling* operation).

**Figure 1:**
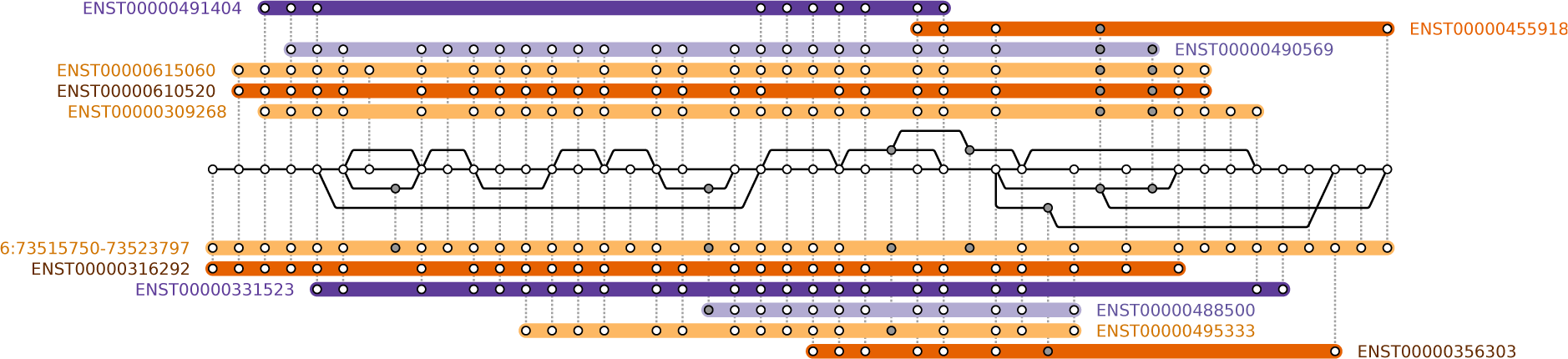
A sequence graph assembled from the *Homo sapiens* reference genome and all transcripts of the EEF1A1 gene in Ensembl v80 [93]. Each node represents a sequence found in the reference genome or transcripts, edges (black solid lines) indicate adjacencies that have been observed in transcripts, and colored horizontal bars correspond to paths of known transcripts (annotated with their Ensembl transcript ID), where each dotted line indicates that a node in the graph is part of this transcript. The path annotated with coordinates (6:73515750– 73523797) corresponds to the reference genome.

#### Comparison among Pan-Genomes

Given any pair of genomes within a pan-genome, we expect a data structure to highlight differences, variable and conserved regions, as well as common syntenic regions. Beyond that, a global comparison of two (or more) pan-genomes, e.g. with respect to gene content or population differentiation, should be supported (see Figure 2, *Compare* operation).

#### Simulation

A pan-genome data structure should support the generation (sampling) of individual genomes similar to the genomes it contains (see Figure 2, *Simulate* operation).

#### Visualization

All information within a data structure should be easily accessible for human eyes by visualization support on different scales (see Figure 2, *Visualize* operation). This includes visualization of global genome structure, structural variants on genome level and local variants on nucleotide level, but also biological features and other computational layers (see paragraph Biological Features and Computational Layers) should be represented.

#### Efficiency

We expect a data structure to use as little space on disk and memory as possible, while being compatible to computational tools with a low running time. Supporting specialized hardware, such as general purpose graphics processing units (GPGPUs) or field-programmable gate arrays (FPGAs), is partly an implementation detail. Yet, in some cases, the target platform can influence data structure design significantly.

### 4.2 Approaches

There are natural trade-offs between some of the desiderata discussed in the previous section. For instance, the capability to allow dynamic updates might be difficult to achieve while using only small space and allowing for efficient indexing. It is one of the core challenges of computational pan-genomics to design data structures that support (some of) the above query types efficiently. While desirable in principle, we consider it difficult, if not impossible, to develop a solution that meets *all* the listed requirements at once. Therefore, future research should aim to delineate the compromises that may have to be made and thereby provide guidance on which solution is suitable for which application scenario. As the field matures, additional queries will appear, and data structures will need to adapt to support them.

In the following, we discuss traditional approaches to meet fundamental requirements for genome analysis, first extensions for pan-genomes, as well as future challenges.

#### Unaligned Sets of Sequences

The conceptually simplest representation of a pan-genome consists of a set of individual sequences (Figure 3a), which might be either whole genomes or parts of it. The traditional view of a species’ pan-genome as the set of all genes [14], which is prevalent in microbiology, can be considered an example for this. Unaligned whole genome sequences on the other hand are, in general, of limited utility for most applications, especially when the genomes are long. So we consider collections of individual genomes mostly as input to build the more advanced representations discussed in the following.

**Figure 2:**
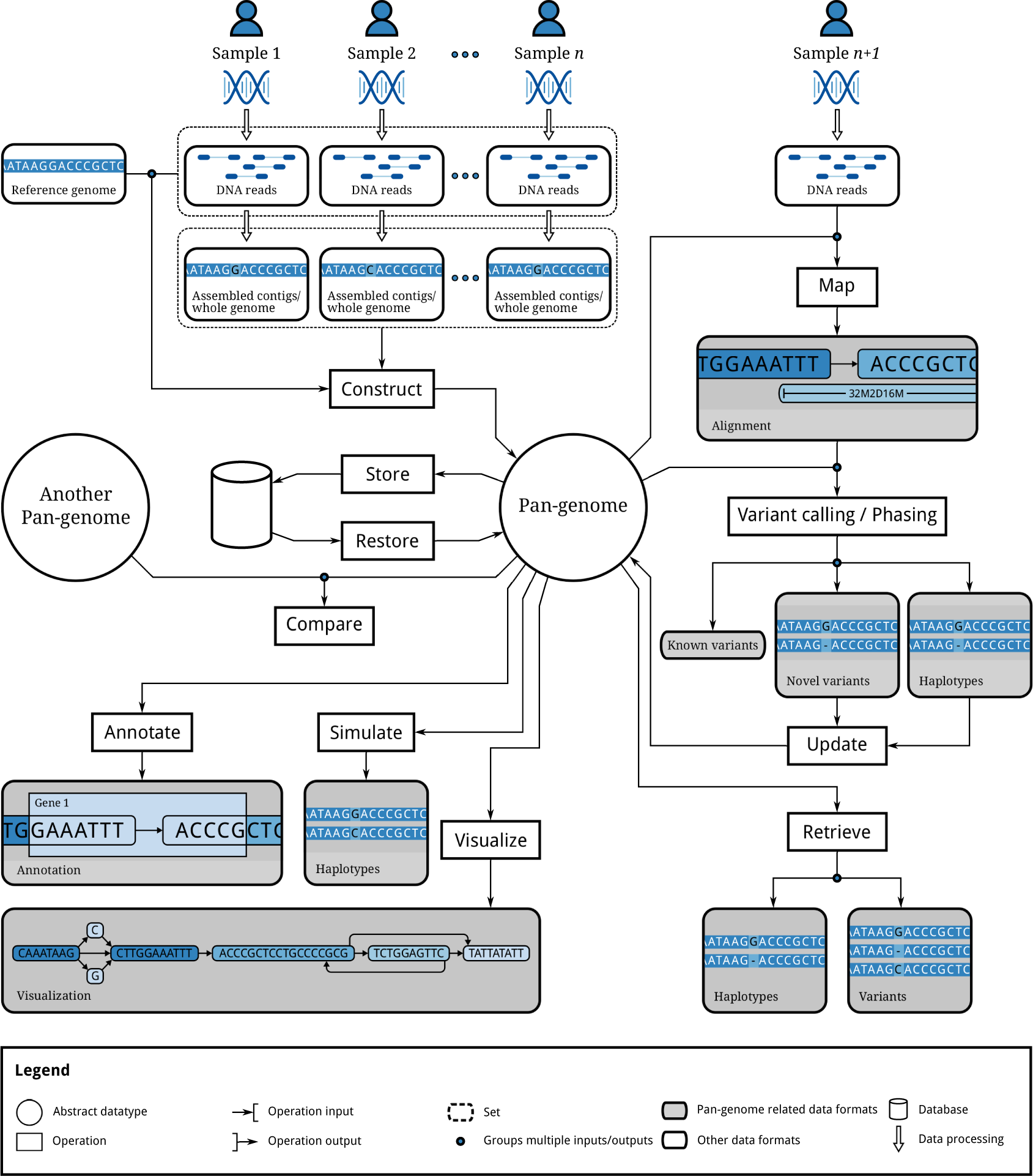
Illustration of operations to be supported by a pan-genome data structure.

#### Multiple-Sequence-Alignment Based Representations

Pan-genomes can be represented by alignments of multiple genomes. In a multiple sequence alignment (MSA), the input sequences are aligned by inserting gap characters into each sequence (Figure 3b). The result is a matrix, where each column represents putatively homologous characters. Refer to [95, 96] for reviews on current methods and remaining challenges. Such classical colinear alignments are not able to capture larger rearrangements like inversions and translocations well and hence only apply to short genomic regions such as single genes or to very similar genomes.

**Figure 3:**
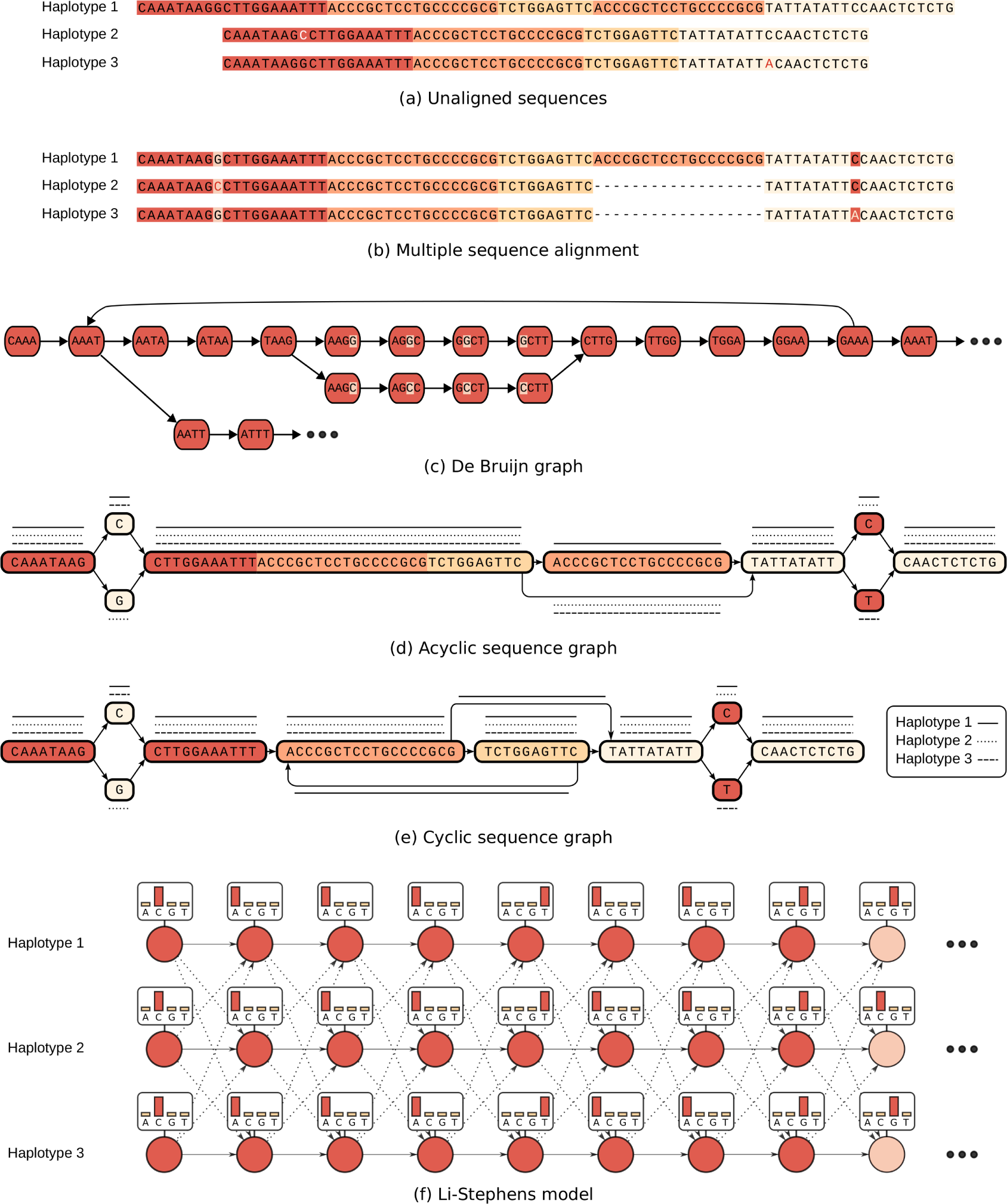
Selected examples of pan-genome representations: (a) three unaligned sequences, colors highlight similarities; (b) a multiple sequence alignment of the same three sequences; (c) the De Bruijn graph of the first (red) sequence block; (d) acyclic sequence graph, paths representing the three haplotypes shown as solid/dashed/dotted lines; (e) cylic sequence graph; (f) Li-Stephens model of the first nine characters with states indicated by circles, emission distributions given in boxes and transitions given by arrows; dashed arrows indicate the (less likely) “recombination” transitions.

One advantage of using an MSA as a representation of a pan-genome is that it immediately defines a coordinate system across genomes: a column in the alignment represents a location in the pangenome. MSAs furthermore support many comparison tasks.

All approaches designed for linear reference genomes can, in principle, be extended to multiple alignments at the expense of adding bookkeeping data structures to record where the gaps are. Efficient data structures for prefix sum, rank, and select queries exist [97], which can be used for the purpose of doing projections to and from a sequence and its gapped version as a row of an MSA. Multiple sequence alignments can be compactly represented by journaled string trees [98]. This data structure also allows for efficiently executing sequential algorithms on all genomes in the MSA simultaneously. One example for such a sequential algorithm is online pattern matching, that is, searching all genomes for the exact or approximate occurrence of a pattern without building an index structure first.

When aligning two or more whole genomes, structural differences such as inversions and translocations need to be taken into account. Standard methods for a colinear MSA are therefore not applicable. Instead, one aims to partition the input genomes into blocks such that sequences within blocks can be aligned colinearly. Creating such a partitioning is a non-trivial task in itself and mostly approached through graph data structures that represent local sequence similarities and adjacencies. On the one hand, such graphs therefore facilitate whole genome alignment. On the other hand, they can be understood as representations of the pan-genome. Concrete realizations of this idea include A-Bruijn graphs [99], Enredo graphs [100] and Cactus graphs [101, 102]. For detailed definitions and a comparison of these concepts we refer the reader to the review [103].

Block-based multiple sequence alignments can also serve as the basis for a coordinate system on a pan-genome: by numbering blocks as well as numbering columns inside each colinearly aligned block, a notion of a position in a pan-genome can be defined. This idea is explored by Herbig et al. [104], who furthermore show how it can serve as a foundation for visualization.

#### k-mer-Based Approaches

Starting from either assembled genomes, contigs, or just collections of (error-corrected) reads, a pan-genome can also be represented as a collection of *k*-mers, i.e. strings of length *k*. The task of efficiently counting all *k-* mers occurring in an input sequence has been studied extensively in recent years and many solutions are available, including Jellyfish [105], DSK [106]and KMC2 [107]. Such a *k*-mer collection is a representation of the corresponding de Bruijn Graph (DBG), illustrated in Figure 3c. DBGs were introduced in the context of sequence assembly [108], but can be used as pan-genome representations supporting many applications beyond assembly. When *k*-mer neighborhood queries are sufficient, and no *k*-mer membership queries are required, then even more space-efficient data structures for DBGs exist [109].

When building DBGs for multiple input samples, one can augment each *k*-mer by the set of samples containing it. This idea is realized in colored DBGs where we color each *k*-mer according to the input samples it occurs in. Colored DBGs have been used successfully for reference-free variant calling and genotyping [110]. Recently, Holley et al. [111]introduced Bloom filter tries, a data structure able to efficiently encode such colored DBGs.

For *k*-mer based representations of pan-genomes, the length *k* is obviously an important parameter and picking the right value depends on the intended application. Data structures able to represent a pan-genome at different granularities (i.e. at different values of *k*) are hence an interesting research topic. For instance, Minkin et al. [112] show that iteratively increasing *k* helps to capture nested synteny structure.

Pan-genomes encompassing many species can be encoded as a mapping between *k*-mers and clades: given a phylogenetic tree, each *k*-mer is mapped to the lowest common ancestor of all genomes containing it. This technique was introduced by Wood and Salzberg [113], who show that it efficiently supports the task of analyzing the composition of metagenomic samples.

Advantages of *k*-mer-based representations include simplicity, speed, and robustness: it is not necessary to produce an assembly or an alignment, which both can be error-prone, and very efficient Bloom-filter-based data structures to store them exist and are available in mature software libraries such as the GATB [114]. However, they do not explicitly represent structural information at distances greater than the *k*-mer length. For applications where such information is needed, DBGs can sometimes serve as a basis to design richer data structures. Colored DBGs [110, 111] are an example of this since they store information about occurrence in individual genomes on top of each *k*-mer.

#### Further Sequence Graphs

Building on the above ideas, more general approaches conceptualize a pan-genome as an (edge-or node-labeled) graph of generic pieces of sequence. Such graphs are not necessarily constructed using an MSA and the constituting sequences are not necessarily fixed-length *k*-mers. Figures 3d and 3e show examples of an acyclic and a cyclic sequence graph, respectively. Individual genomes can be represented as paths in such graphs and node identifiers can serve as a “co-ordinate system”.

Compressed DBGs (also called compacted DBGs), which collapse chains of non-branching nodes in a DBG into a single node, are an example of this. Marcus et al. [115] show how such compressed DBGs can be constructed for a pan-genome by first identifying maximal exact matches using a suffix tree, by-passing uncompressed DBGs. Beller and Ohlebusch [116] and Baier et al. [117] show how the same can be achieved more efficiently, using an FM index resp. compressed suffix trees and the Burrows-Wheeler transform. Trading efficiency for comprehensibility, compressed DBGs can also form the foundation for annotated pan-genomes stored in graph databases [?].

Useful data structures for pan-genomes may combine some of the basic approaches discussed so far. For example, PanCake [118] uses a graph-based structure to represent common genomic segments and uses a compressed multiple-alignment based representation in each node of the graph. Dilthey et al. [15] propose a generative model by representing sequence variation in a *k*-mer-emitting HMM.

Further examples of implementations of sequence graphs include the Global Alliance for Genomics and Health (GA4GH) “side graph” data model and the FASTG format^6^. Side graphs represent a pangenome as a set of sequences and an additional set of joins, each of which defines an extra adjacency between the sides of two bases within the sequences. The GA4GH graph tools^7^ allow side graphs and embeddings of individual sampled genomes in that graph to be made available over the Internet, for data distribution and remote analysis.

#### Haplotype-Centric Models

When a fixed set of (non-nested) sequence variants is considered, every haplotype in a population can be represented as a string of fixed length. The character at position *k* reflects the status of the *k*-th variant. When all variants are bi-allelic, then these haplotype strings are formed over a binary alphabet. Such collections of haplotypes are often referred to as *haplotype panels*. This representation is favorable for many population genetic analyses since it makes shared blocks of haplotypes more easily accessible, for instance compared to sets of paths in a graph.

A recent data structure to represent haplotype panels, termed Positional Burrows-Wheeler Transform (PBWT) [119], facilitates compression and supports the enumeration of maximal haplotype matches.

One of the most widely used haplotype-based models is the Li-Stephens model [120]. In a nutshell, it can be viewed as a hidden Markov model (HMM) with a grid of states with one row per haplotype and one column per variant, as sketched in Figure 3f. Transitions are designed in a way such that staying on the same haplotype is likely but jumping to another one is also possible. It hence is a generative probabilistic model for haplotypes that allows for sampling new individuals and provides conditional probabilities for new haplotypes given the haplotypes contained in the model.

## 5 Computational Challenges

Pan-genomic data have all of the standard properties of *Big Data*—in particular, volume, variety, velocity and veracity. Especially due to the sheer size of generated sequencing data, extreme heterogeneity of data and complex interaction on different levels, pan-genomics comes with big challenges for algorithm and software development [121]. The International Cancer Genome Consortium (ICGC) has amassed a dataset in excess of two petabytes in just five years with the conclusion to store data generally in clouds, providing an elastic, dynamic and parallel way of processing data in a cheap, flexible, reliable and secure manner [122].

Currently large computing infrastructure providers and large public repositories (e.g. NCBI/EBI/DDBJ) are completely separated. We need hybrids that offer both large public repositories as well as the computing power to analyze these in the context of individual samples/data. We consider it desirable to bring the computation as close as possible to the data by uploading queries or in-database computing.

*Distributed and parallel computing* will be necessary to successfully handle pan-genome data in practice. To this end, leveraging the capabilities of existing Big-Data frameworks is desirable and should be combined with bringing the computation as close as possible to the data. On the practical side, tackling these challenges will also involve establishing widely-accepted standards for *file formats* for sequence graphs and related data like read alignments to such graphs. On the theoretical side, studying changes in algorithmic complexity when working with sequence graphs instead of sequences will be an interesting and challenging aspect of computational pan-genomics.

All these general challenges apply to all individual computational problems we discuss in the following.

### 5.1 Read Mapping

Given a set of reads sequenced from a donor, *read mapping* consists in identifying parts of the reference genome matching each read. Read mapping to a pan-genome has a potential to improve alignment accuracy and subsequent variant calling, especially in genomic regions with a high density of (complex) variants.

For a single reference sequence, the read mapping problem has mostly been solved by indexing the reference into a data structure that supports efficient pattern search queries. Most successful approaches use *k*-mer based or Burrows-Wheeler transform based indexes, as reviewed in [123]. Indexing a pan-genome is more complicated.

Efficient indexing of a set of reference genomes for read mapping was first studied in [124, 125]. The approach uses compressed data structures, exploiting the redundancy of long runs of the same letter in the Burrows-Wheeler transform of a collection of similar genomes. This approach yields a reasonably compressed representation of the pangenome, but read alignment efficiency is hampered by the fact that most reads map to all of the references, and that extraction of these occurrence locations from a compressed index is potentially slow. More recently, approaches based on LempelZiv compression have been proposed to speed-up the reporting of occurrences, as reviewed in [126].

The earliest approach to index a *sequence graph* (see Section 4. 2) was proposed in [127], where *k-* mer indexing on the paths of such a graph was used; instead of a full sequence graph, a *core* sequence graph was used where columns were merged in regions of high similarity (core genome) to avoid extensive branching in the graph. After finding seed occurrences for a read in this graph, the alignment was refined locally using dynamic programming. Similar *k*-mer indexing on sequence graphs has since been used and extended in several read mapping tools such as MuGI [128], BGREAT [129] and VG^8^.

Instead of *k*-mer indexing, one can also use Burrows-Wheeler-based approaches, based on appending extracted contexts around variations to the reference genome [130]. Context extraction approaches work only on limited pattern length, as with long patterns they suffer from a combinatorial explosion in regions with many variants; the same can happen with a full sequence graph when all nearby *k*-mer hit combinations are checked using dynamic programming. There is also a special Burrows-Wheeler transform and an index based on that for a sequence graph [131, 132]. This approach works on any pattern length, but the index itself can be of exponential size in the worst case; best case and average case bounds are similar to the run-length compressed indexes for sets of references like [125]. The approach is also likely to work without exponential growth on a core sequence graph of [127], but as far as we know, this combination has not been explored in practice. A recent implementation^9^ avoids the worst case exponential behavior by stopping the construction early; if this happens, the approach also limits the maximum read length. This implementation has been integrated into VG as an alternative indexing approach. HISAT2^10^ [94] implements an index structure that is also based on [131], but builds many small index structures that together cover the whole genome.

In summary, a number of approaches to perform read mapping against a pan-genome reference under various representation models exist, and efficient implementations for daily usage are under active development. However, we consider this field as being far from saturated and still expect considerable progress in both algorithmic and software engineering aspects. To reach the full potential of these developments, the interactions between read mapping and variant calling methods need to be considered.

### 5.2 Variant Calling and Genotyping

The task of determining the differences between a sequenced donor genome and a given (linear) reference genome is commonly referred to as *variant calling*. In case of diploid or polyploid organisms, we additionally want to determine the corresponding *genotype*. In the face of pan-genome data structures, variant calling decomposes into two steps: identifying *known* variants already represented in the data structure and calling *novel* variants. Refer to Schneeberger et al. [127] for an early work on pan-genome variant calling. They do not only show the feasibility of short read alignment against a graph representing a pan-genome reference (see Section 5. 1) but also demonstrate its positive impact on variation calling in the frame of the Arabidopsis 1001 Genomes Project.

#### Known Variants

By using a pan-genome reference, one merges read mapping and calling of known variants into a single step. Read alignments to sequence variants encapsulated in our pan-genome data structure indicate the presence of these variants in the donor genome. In particular, this applies not only to small variants which can be covered by a single read (such as SNPs and indels), but also to larger structural variants such as inversions or large deletions. Integrating those steps potentially decreases overall processing time and, more importantly, removes read-mapping biases towards the reference allele and hence improves accuracy of calling known variants. One important challenge is to statistically control read mapping ambiguity on a pan-genome data structure. Leveraging the associated statistical models for estimating genotype likelihoods is expected to lead to significant improvements in genotyping.

As a first major step in that direction, Dilthey et al. [15] cast the (diploid) variant calling problem into finding a pair of paths through a pangenome reference represented as a *k*-mer-emitting Hidden Markov Model. They demonstrate that this leads to substantially improved performance in the variation-rich MHC region.

#### Novel Variants

Detecting variants not present in a pan-genome data structure is similar to traditional variant calling with respect to a linear reference genome. Still, differences exist that require special attention. The most straightforward way to use established variant calling methods is to use read alignments to a pan-genome and project them onto a linear sequence. For small variants such as SNPs and indels, that are contained within a read, this approach is likely to be successful. Methods to characterize larger structural variation (SV) need to be significantly updated. SV calling methods are usually classified into four categories based on the used signal: read pair, read depth, split read, and assembly, as reviewed by Alkan et al. [59]. Each of these paradigms has its merits and shortcomings and state-of-the-art approaches usually combine multiple techniques [133]. Each of these ideas can and should be translated into the realm of pangenomes. For split-read and assembly based approaches, the problem of aligning reads and contigs, respectively, to a pan-genome data structure (while allowing alignments to cross SV breakpoints) needs to be addressed. In case of read pair methods, a different notion of “distance” is implied by the pan-genome model and has to be taken into account. For read depth methods, statistical models of read mapping uncertainty on pan-genomes have to be combined with models for coverage (biases). Developing standards for reporting and exchanging sets of potentially nested variant calls is of great importance.

#### Somatic Mutations

Calling somatic mutations from paired tumor/normal samples is an important step in molecular oncology studies. Refer to Section 2.6 for details and to [134] for a comparison of current work flows. Calling somatic variants is significantly more difficult compared to calling germ-line variants, mostly due to tumor heterogeneity, the prevalence of structural variants, and the fact that most somatic variants will be novel. Pan-genome data structures promise to be extremely useful in cancer studies for the stable detection of somatic variants. A conceivable approach for leveraging pan-genome data structures in this context would be to map reads from the matched normal sample to the pan-reference, call germline mutations, create a restricted pan-genome with detected variants and map tumor reads to that pan-reference for calling somatic mutations. There are many more potential applications including building a pan-genome representation of a heterogeneous tumor to be used as a starting point for retracing tumor evolution.

#### Storing Variants

Storing and exchanging variant calls genotyped in a large cohort of samples increasingly becomes a bottleneck with growing cohort sizes. Some improvement is achieved by adopting binary instead of text-based data formats for variant calls, i.e. using BCF instead of VCF^11^, but more efficient approaches are urgently needed. Organizing data by individual rather than by variant while sorting variants by allele frequency has proven beneficial for compression and some retrieval tasks [135]. We expect the question of storing, querying and exchanging variant data to remain an active and relevant field of research in the coming years.

### 5.3 Haplotype Phasing

Humans are diploid, that is, each chromosome comes in two copies, one inherited from the mother and one inherited from the father. The individual sequences of these two chromosomal copies are referred to as *haplotypes*, where one often restricts the attention to polymorphic sites. The process of assigning each allele at heterozygous loci to one of the two haplotypes is referred to as *phasing*. Plants are often polyploid. For example, wheat can be tetra(= 4 copies) or hexaploid (= 6 copies), while certain strawberries are even decaploid (= 10 copies). As an extreme, the “ploidy” of viral quasispecies, that is the number of different viral strains that populate an infected person (see Section 2. 3) is usually unknown and large. The same applies to heterogeneous tumors, as discussed in Section 2. 6.

Pan-genome data structures have the potential to, on the one hand, store haplotype information and, on the other hand, be instrumental for phasing. Currently, several approaches for obtaining haplotype information exist, as reviewed in [79, 136]. *Statistical phasing* [137] uses genotype information of large cohorts to reconstruct haplotypes of all individuals based on the assumption that haplotype blocks are conserved in a population. Once sets of haplotypes, called reference panels, are known, additional individuals can be phased by expressing the new haplotypes as a mosaic of the already known ones. The question of how to best organize and store reference panels is open. To this end, Durbin [119] has proposed the aforementioned PBWT index structure. We consider marrying reference panels to pan-genome data structures an important topic for future research.

To determine haplotypes of single individuals, including rare and *de novo* variants, statistical approaches are not suitable and experimental techniques to measure linkage are needed. Such techniques include specialized protocols and emerging long-read sequencing platforms, as discussed in Section 3. Currently, first approaches for haplotype-resolved local assembly are being developed [138]. More literature exists on the problem of phasing from aligned long reads, e.g. [139, 140, 141]. In practice, this technique is hampered by insufficient alignment quality of long error-prone reads. Since phasing is based on heterozygous loci, avoiding allelic biases during read mapping by means of pangenome data structures can contribute to solving this problem. Combining the virtues of read-based phasing with statistical information from reference panels is an active area of research [87]. Leveraging pan-genome data structures that encode reference haplotypes towards this goal constitutes a promising research direction.

These problems are amplified when phasing organisms or mixtures of higher or unknown ploidy such as plants, viral quasispecies or tumors. Algorithms with manageable runtime on polyploid organisms [142, 143] and for the reconstruction of quasispecies [144, 145] require the use of specialized techniques (especially when allele frequencies drop below sequencing error rates). Extending these approaches to pan-genome data structures is another challenging topic for future research.

## 5.4 Visualization

Pan-genomics introduces new challenges for data visualization. Fundamentally, the problems relate to how to usefully view a large set of genomes and their homology relationships, and involve questions of scale and useful presentation in the face of huge volumes of information.

At a high-level of abstraction, pan-genome bagof-genes approaches can be visualized using methods for comparing sets, such as Venn diagrams, flower plots, and related representations. For example, the recent tool Pan-Tetris visualizes a gene-based pan-genome in a grid [146], color-coding additional annotation. For divergent genomes, as in bacterial- and meta- pan-genomics, and where complete assembly is not possible, such approaches provide useful summary information.

For the viewing of individual, assembled genomes or sequences, genome browsers and applications frequently display an individual sequence along a linear or circular axis upon which other genomics information is visualized, as reviewed in [147]. This trope, which is popular and widely understood, forces interpretation through the lens of one chosen genome. When this genome is a distantly related reference genome there is a visual reference bias which may lead to misinterpretation.

Pan-genome displays can potentially help to alleviate this visual bias. One option is to aim to improve linear visualizations: either the chosen individual reference sequence can be replaced by a more visually useful imputed pan-genome reference, or the pan-genome data structures which relate different genomes in the population can be used to translate information to the most closely related genome possible for the display. In the former case, a pan-genome display can be made more inclusive than any single genome [148]. At the base level such inclusive displays are somewhat analogous to popular multiple sequence alignment displays such as Mauve [149] or Jalview [150] that focus on displaying all the differences between a set of sequences as clearly as possible. The latter case, translation, where a pan-genome alignment is used to show information on the most closely related genome possible, is likely to become more popular as the number of available personal genomes grows, see [25] for an early example of such an approach.

More adventurously than linear layouts, pangenome displays can attempt to visualize graphs of variation. This has the flexibility of allowing arbitrary genome variation within a clean semantic model, but can prove visually complex for even small, non-trivial examples. For example, a graph of a few dozen bacterial strains contains tens to hundreds of thousands of nodes and edges. So far graph visualizations have proved popular for assemblies, and the visualization of heterozygosity, for example DISCOVAR [151] contains a module that allows you to visualize subsets of an assembly graph in a figure. One popular tool is Cytoscape [152], which is a generic biological graph/network visualization tool, but lacks scalability and semantic navigation. Another tool, Bandage [153], visualizes *de novo* assembly graphs specifically.

A number of challenges exist moving forwards. In a useful visualization it will be possible to navigate and to zoom in and out on pan-genome structures. Zooming should be done semantically, i.e. different zoom levels can use different representations of the data to convey biologically relevant information. The upper scales should give information about global genome structure. Zooming in the visuals should focus on structural variants in a genomic region and the most zoomed in views should enable exploration of local variants on nucleotide level. Furthermore these visuals need to be put in the context of the phylogeny, e.g. the relation of the various samples that went into the pan-genome. This will enable rapid identification and interpretation of observed variants. Finally, any pan-genome graph visualization should offer the same basic features that current reference based genome browsers have. There should be visual ways to indicate biologically interesting features such as gene annotations and position based continuous valued signals such as wiggle tracks in the UCSC genome browser. Basic analytical capabilities would be beneficial to visually highlight interesting biologically relevant mutations. For example, it would be useful to have different visual representations for different types of mutations: indels, (non)-synonymous SNPs, structural variants, repeats etc.

## 5.5 Data Uncertainty Propagation

One of the computational (and modeling) challenges facing the field of pan-genomics is how to deal with data uncertainty propagation through the individual steps of analysis pipelines. In order to do so, the individual processing steps need to be able to take uncertain data as input and to provide a ‘level of confidence’ for the output made. This can, for instance, be done in the form of posterior probabilities. Examples where this is already common practice include read mapping qualities [154]and genotype likelihoods [155].

Computing a reasonable confidence level commonly relies on weighing alternative explanations for the observed data. In the case of read mapping for example, having an extensive list of alternative mapping locations aids in estimating the probability of the alignment being correct. A pan-genome expands the space of possible explanations and can, therefore, facilitate the construction of fairer and more informative confidence levels.

As an illustration, consider a pipeline including read mapping, variant calling and genotyping, phasing and association testing. Substantial uncertainty and sequence composition biases are already inherent to the input data generated by next-generation sequencing [156]. The following read alignment step adds ambiguity in read placement, leading to uncertain coverage and fragment lengths. As a result, this leads to uncertainties in variant calling, genotyping, and phasing. This, finally, results in uncertainties in association testing in genome-wide association studies. The precise quantification of the propagation of these effects is largely unclear. The advent of ever larger and refined panels, supported by appropriate pan-genome data structures, bears the promise of making quantification and alleviation of such effects possible.

## 6 Conclusions

Already today, the DNA having been sequenced for many biologically coherent ensembles—such as certain taxonomic units or virus populations—likely captures the majority of their frequently occurring genetic variation. Still, the pace at which genomes are currently sequenced is on a steep rise, thanks to accumulation of sequencers in laboratories and frequent, significant advances in sequencing technology. Therefore, capturing *all of genomes*, in terms of genetic variation content and abundance, is no longer wishful thinking, but will materialize soon for many species, populations, and cancer genomes. In other words, life sciences have entered the era of *pan-genomics*, which is characterized by knowing *all* major genetic variation of a collection of genomes of interest. In this paper, we addressed the question of how to arrange and analyze this incredible wealth of knowledge and also how to deal with some of the consequences in downstream analyses.

### 6.1 Present Status

The computational aspects that need to be considered fan out across a large variety of particular challenges, usually governed by the realm of application they stem from. We have listed the many facets of pan-genomes in terms of functionality, annotational detail, computational efficiency issues and visualization. We have discussed how the availability of well-arranged pan-genomes will affect population genetics, cancer genomics, pathogen research, plant breeding, phylogenomics, functional genomics as well as genetic disease research and genome-wide association studies. We have surveyed the impact of sequencing technology advances on the field of pan-genomics, and we have considered also the complications that come along with these advances. We have put particular emphasis on data structures and supporting algorithms that make it possible to consistently work with pan-genomes. One of the currently most evident processes in computational pan-genome research is the move away from linear reference genomes towards reference systems that are rooted in graph theory in some form. The effort of the Data Working Group of the Global Alliance for Genomics and Health (GA4GH) is a prominent example for this. We have also discussed how the transition in terms of data structures will affect operations such as read mapping, variant discovery, genotyping and phasing, all of which are at the core of modern genomics research. Last but not least, we have analyzed the issues that arise in visualizing pan-genomes, and we have also briefly discussed future issues in uncertain data handling, recently an ever recurring theme in genome data analysis, often arising from the repetitive structure of many genomes.

We have concentrated on computational challenges of pan-genomics in this survey. We are aware that there are also political challenges that have to be addressed that concern data sharing and privacy. Clearly, the usefulness of any pan-genomic representation will increase with the number of genomes it represents, strengthening its expressive and statistical power. Unfortunately, however, only a fraction of the sequenced data is currently publicly available. This is partly due to the confidential nature of human genetic data, but also, to a large extent, by missing policies and incentives to make genomic data open access or to prevent intentional withholding of data. Funding agencies like the National Institutes of Health (NIH) in the US have started to address these issues [157]^12^.

### 6.2 Future Directions

Overall, we have provided a broad overview of computational pan-genomics issues, which we hope will serve as a reference for future research proposals and projects. However, so far, we have mostly been addressing how to deal with genomes as sequences, that is from a “one-dimensional” point of view, and so we have been focusing on storing and analyzing sequences and the mutual relations of particular subsequence patches, like variant alleles and their interlinkage, genes and/or transcriptomes. We have done this because we believe that at this point in genomics history, only the consistent exploration and annotation of exhaustive amounts of sequence information can lay the solid foundation for additional “pan-genomics oriented” steps.

Yet, even after having resolved the corresponding issues—and we are hopeful that, at this point, our summary has helped to consistently structure these—there is more to follow. New approaches have already appeared on the horizon that will benefit from the cornerstone provided by primarily sequence-driven pan-genomics. For example, it can be expected that one can lift pan-genomes into three dimensions in the mid-term future, thanks to rapidly developing techonolgies that allow to infer their three-dimensional conformation. This will mean that future, three-dimensional pan-genomes will not only represent all sequence variation applying for species or populations, but also encode their spatial organization as well as their mutual relationships in that respect.

Epigenomics topics have not been exhaustively addressed here either, but will need to be addressed as soon as the first “primary” pan-genomes stand. Technologies that do not only map sequential and three-dimensional arrangement, but also additional biochemical modifications have likewise been on a steep rise recently. Most importantly, we will be in position to link sequential pan-genomes to maps that indicate hypo- and hypermethylated regions relatively soon. Likely, the integration of such basic biochemical modifications will serve as template for further, often more complex elements of biochemical genomic maps.

In summary, the emergence of computational pan-genomics as a field is a major advance in contemporary genomics research. We have entered an era that holds the promise to close large gaps in global maps of genomes and to draw the full picture of their variability. We therefore believe that we can expect to witness amazing, encompassing insights about extent, pace, and nature of evolution in the mid-term future.

## Acknowledgments

We are deeply grateful to the Lorentz Center for hosting the workshop “Future Perspectives in Computational Pan-Genomics” (June 8–12, 2015), which gave rise to this paper. In particular, we like to thank the Lorentz Center staff, who turned organizing and attending the workshop into a great pleasure. The workshop received additional financial support by KNAW, Bina Technologies, ERIBA, PacBio, and Genalice.

## Competing Interests

Ole Schulz-Trieglaff is an employee of Illumina Inc. and receives stocks as part of his compensation. Illumina is a public company that develops and markets systems for genetic analysis.

#### Key points

- Many disciplines, from human genetics and oncology to plant breeding, microbiology and virology, commonly face the challenge of analyzing rapidly increasing numbers of genomes.
- Simply scaling up established bioinformatics pipelines will not be sufficient for leveraging the full potential of such rich genomic datasets.
- Novel, qualitatively different computational methods and paradigms are needed and we will witness the rapid extension of computational pan-genomics, a new sub-area of research in computational biology.
- The transition from the representation of reference genomes as strings to representations as graphs is a prominent example for such a computational paradigm shift.

## Footnotes

1 See http://www.technologyreview.com/news/537916/rebooting-the-human-genome, for an example of recent media coverage

2 http://irri.org/our-work/research

3 http://omim.org/

4 http://evs.gs.washington.edu/EVS

5 http://exac.broadinstitute.org

6 http://fastg.sourceforge.net

7 https://github.com/ga4gh/server and https://github.com/ga4gh/schemas

8 https://github.com/ekg/vg

9 https://github.com/jltsiren/gcsa2

10 https://ccb.jhu.edu/software/hisat2/index.shtml

11 http://samtools.github.io/hts-specs/

12 see also http://www.nih.gov/news-events/news-releases/nih-issues-finalized-policy-genomic-data-sharing

